# Model-free prediction of microbiome compositions

**DOI:** 10.1101/2022.02.04.479107

**Authors:** Eitan E. Asher, Amir Bashan

## Abstract

The recent recognition of the importance of the microbiome to the host’s health and well-being, has yielded efforts to develop therapies that aim to shift the microbiome from a disease-associated steady-state to a healthier one. Direct manipulation techniques of the species’ assemblage are currently available, e.g., using probiotics or narrow-spectrum antibiotics to introduce or eliminate specific taxa. However, predicting the species’ abundances at the new steady-state remains a challenge, mainly due to the difficulties of deciphering the delicate underlying network of ecological interactions or constructing a predictive model for such complex ecosystems. Here, we propose a model-free method to predict the species’ abundances at the new steady state based on their presence/absence configuration by utilizing a multi-dimensional k-nearest-neighbors (kNN) regression algorithm. By analyzing data from numeric simulations of ecological dynamics, we show that our predictions, which consider the presence/absence of all species holistically, outperform both the null model that uses the statistics of each species independently and a predictive neural network model. We analyze real metagenomic data of human-associated microbial communities and found that by relying on a small number of ‘neighboring’ samples, i.e., samples with similar species assemblage, the kNN predicts the species abundance better than the whole-cohort average. By studying both real metagenomic and simulated data, we show that the predictability of our method is tightly related to the dissimilarity-overlap relationship of the training data. Our results demonstrate how model-free methods can prove useful in predicting microbial communities and may facilitate the development of microbial-based therapies.

## Introduction

Motivated by the significant role that microbial communities play in various environments [1–3], ongoing efforts are being made to develop practical therapies for diseased human-associated microbial communities and rehabilitation techniques for disrupted natural microbial ecosystems [4, 5]. The successful development of rational and safe manipulations of microbial communities depends on the ability to predict their post-perturbative composition. In practice, typical intervention techniques for microbial communities involve manipulation of the ‘species assemblage’, i.e., the sample-specific configuration of resident species. Such manipulations include removal of microbial species, e.g., using narrow or broad-spectrum antibiotics, or introduction of single cultured probiotic organisms, consortia of microbes, or ‘complete’ microbial ecologies, e.g., fecal microbiota transplants. After such perturbation, the modified ecosystem is expected to be shifted towards a new steady-state where the species abundances are determined by the new ecological balance. Therefore, we aim to predict the abundance composition of microbial communities based on their species assemblage.

Current methods to predict the composition of microbial communities commonly rely on prior reconstruction of a mediating network model. Such approaches mainly describe an effective population dynamics model whose parameters are chosen by fitting to the available data, or from detailed knowledge of the biochemical reactions [6–9]. These models predict the personalized response of a particular microbial community to a given perturbation by tracking the resultant time-dependent dynamics. However, applying such models to large real-world microbial communities is very challenging, mainly due to the difficulty in reliably fitting their parameters [9, 10], and concerns regarding their validity [11].

Another active direction is to apply machine learning techniques to microbial communities [12–21]. These methods have proven useful in various tasks, including in predicting the steady state of microbial communities using deep learning [22]. However, training a predictive model has several caveats. For example, it often requires a large number of training samples as well as extensive computational and time resources. In addition, the trained model might not be directly generalized to different datasets or different ecological communities.

Here, we propose a model-free approach to predict the composition of microbial communities based on their species assemblage. We assume that in the analyzed ecological dynamics there is a one-to-one mapping, relating each species assemblage to a unique steady-state composition, i.e., true multi-stability does not exist, but the alternative steady states are associated with different species assemblages. This assumption is mathematically valid for the case of the generalized Lotka Volterra (GLV) model of ecological dynamics and, as far as we know, true multi-stability in human-associated microbial communities has not been demonstrated experimentally.

Our approach avoids the inherent issues involved with the prerequisite of network reconstruction. In addition, it directly predicts the steady state, avoiding the sensitive task of following time-dependent dynamics. This approach is relevant to a broad range of medical conditions for which potential microbiome-based therapies are tested. For example, when developing microbiome manipulations to treat chronic diseases, such as IBD or diabetes, we are mainly interested in the long-term stability of the system rather than its time transition.

## Methodology

In this study, we develop a framework to predict the abundance profile of a test sample **Φ**, based on its species assemblage *φ*. The species assemblage of the test sample is a species subset from a pool of *N* species. Here, **Φ**_*i*_ (*i* = 1 … *N*) is the relative abundance of species *i*, and ϖ_*i*_ is the presence/absence of species *i*, such that ϖ_*i*_ = 0 if Φ_*i*_ = 0 and ϖ_*i*_ = 1 otherwise. The prediction is based on a given “training set” of *m* other abundance profiles, represented as an *m ×N* matrix Θ, i.e., the matrix element Θ_*j,i*_ represents the abundance of species *i* in training sample *j*. The key idea is that we distinguish between two different features in each training sample, its species assemblage and its abundance profile. We utilize a multi-dimensional *k*-nearest neighbors (kNN) regression algorithm where the features are the species assemblages and the target is the abundance profile. Essentially, our method predicts the abundance profile of the test sample as the average profile of the *k* training samples with the species assemblages that are the most similar to the test sample (as illustrated in Fig. 1).

**Figure 1.**
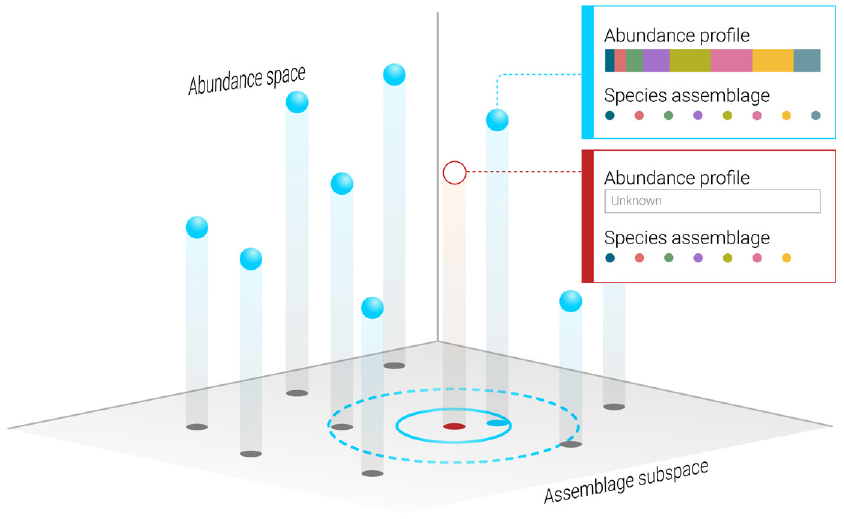
Prediction of microbiome composition based on its species assemblage. An illustration of the framework that is used in order to predict the abundance profile (open red circle), based on its known species assemblage, given a training set of other abundance profiles for which both the species assemblages and their abundances are known (blue spheres). We consider the species assemblages of the training samples, illustrated by ‘projecting’ them onto the assemblage subspace, and the abundance of the sample of interest is predicted based on the samples with the most similar species assemblages, using the k-nearest neighbors (kNN) method.

The specific steps are subsequently elaborated. We start by transforming the training set Θ, into a binary matrix θ of the same size that represents the species assemblages of the training set such that θ_*j,i*_ = 0 if Θ_*j,i*_ = 0 and θ_*j,i*_ = 1 otherwise. Next, we calculate the Jaccard similarity between the species assemblage of the test sample, ***φ***, and the species assemblage of each of the training samples *θ*_*j*_ (*j* = 1 … *m*) (see Methods). The subset of the *k* samples with the highest values of Jaccard similarity, i.e., the *k* nearest neighbors, is denoted as *R*. The average abundance profile of the *k* nearest neighbors 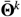 is defined as the average abundance, while the average is calculated over the present species only. Specifically, the average abundance of species *i* is

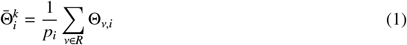

where *p*_*i*_ is the number of samples in *R* for which the abundance of species *i* is non-zero. Finally, the predicted abundance profile of the test sample **Φ** is defined 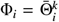 where Φ_*i*_ = 1 and Φ_*i*_ = 0 where Φ_*i*_ = 0. This means that the abundances of the present species in the test sample are predicted according to the average profile of the *k* nearest neighbors, while the abundances of the absent species in the test sample are set to zero. In case *p*_*i*_ = 0 and *φ*_*i*_ = 1, i.e., species *i* is absent in all the *k* nearest neighbors but is present in the test sample, its abundance is predicted as the average abundance over all samples.

To assess the added value of the kNN method, its predictions are compared against an alternative ‘naïve’ approach that predicts the abundance of each present species as the average abundance of that species, i.e., the abundance of species *i* is 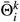 as defined in Eq. 1 if *ϕ*_*i*_ = 1 (species *i* is present in the test sample) and 0 otherwise. This ‘null model’ that predicts each species independently represents an alternative approach to the kNN method which assumes that the abundance predictions are better when taking into account the configuration of all species.

This ‘null model’ represents an alternative approach to the kNN method since it predicts each species independently. In contrast, the kNN method assumes that the abundance predictions are better when taking into account the configuration of all species. In other words, in order to predict the abundance of species *i*, the kNN first considers the species configuration of all other species to select the nearest neighbors, based upon which the abundance of species *i* will be predicted.

## Results

We systematically study the kNN method using simulated and real abundance profiles. We predict the abundance profiles of ‘test samples’ using their species assemblage only and compare them to the real abundance profiles, defining the prediction error as the root Jensen-Shannon divergence (rJSD) between them (see Methods).

To study the effect of the number of training samples on the performance of the kNN method, we test it on synthetic data generated using the GLV model, which has been used for qualitative modelling of the ecosystems [23, 24] (see Methods). We model ecosystems with *N* species (*N* = 10, 20, 40, 100) and calculate the prediction errors when using different sizes of training sets as well as different values of *k* (Fig. 2**a**-**d**). In all cases, the prediction error is reduced for larger sizes of the training set. By increasing the number of training samples, the density of the *N*-dimensional state space increases, resulting in more similar neighbors and better predictions. In marked contrast, the prediction error of the null model remains the same even when large training sets are used.

**Figure 2.**
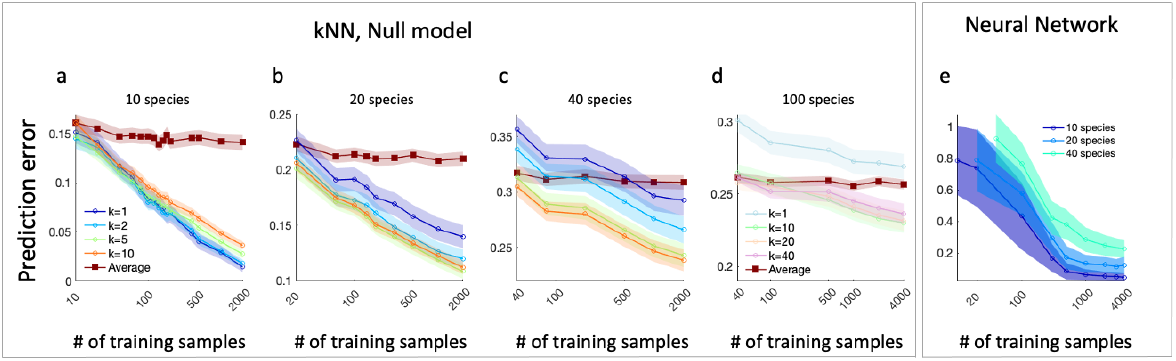
Enhanced prediction accuracy in the kNN method with increasing training samples. We used the GLV model to generate abundance profiles as a training set and 20 test samples, whose abundance profiles were predicted based on their species assemblages. The prediction error is calculated as the rJSD between the predicted and real abundance profiles. Symbols and shaded areas represent the average over 100 realizations and the standard deviation, respectively. **a**, Prediction error of the kNN (circles) and the null model (squares) versus the number of training samples, for a system of 10 interacting species. **b** and **c**, same as **a** for 20 and 40 species, respectively. The kNN predictions improve for a larger number of training samples, in marked contrast with the null model which is effectively independent of the number of training samples. **d** same as **a** for 100 species, here we set *σ* = 0.4. **e**, Results of predictions of a neural network model trained and tested with similar conditions as in **a-c**. Compared with the kNN, the neural network model yields a higher prediction error for a small number of training samples (< 500).

We demonstrate the effect of the number of training samples on the kNN method by comparing it to a neural network model (see Methods), as shown in Fig. 2**d**. While the predictions of the neural network model improve for larger training sets, it requires much more training samples to achieve the same prediction error as the kNN. For example, when using a training set of one hundred samples, the prediction errors of the kNN are about 0.09, 0.18, and 0.28 for 10, 20 and 40 species, respectively, compared with 0.45, 0.6, and 0.8, when using the neural network model (the results of the neural network model are not compared with the case of *N* = 100 which was simulated with weaker interactions to ensure system’s stability). Additionally, the neural network training is considerably time-consuming compared with the kNN, which does not require any training. Obviously, machine learning methods can be applied using a variety of models that might work better in this case. Specifically, we note that the machine learning models were trained using standard algorithms that implement the mean square error loss function. Yet, we consider the studied model as a representative example of the fundamental differences between the kNN as a model-free method and the neural network approach.

An important feature of the kNN method is demonstrated when comparing the effect of *k* (*k* = 1, 2, 5, 10) in the three different system sizes. For the smaller systems (*N* = 10 species) the predictions are the best for *k* = 1, while for larger systems (*N* = 20, 40), the predictions improve for larger values of *k*. Generally, the optimal *k* used in the kNN method balances two effects: i) predictions with larger values of *k* rely on more training samples which improve the statistics, ii) the closer training samples on the features-space represent more relevant and specific information for the particular test sample. In the case shown in Fig. 2**a** representing a system with a pool of 10 species, a typical sample consists of about 6 species and the typical 2nd nearest neighbor has a considerably different species assemblage compared with the test sample. Therefore, for small systems, *k* = 1 produces the best predictions. In contrast, in larger systems (*N* = 20, 40) the prediction errors with *k* = 5, 10 are smaller than *k* = 1, 2, since the 5th or 10th nearest samples still represent a close neighborhood of the test sample.

To test this effect systematically in large and complex ecosystems, we analyze cross-sectional human-associated microbial samples from different body sites from a large-scale metagenomic study, the Human Microbial Project (HMP) [1]. The dataset contains abundance profiles collected from 13 different human body sites, where ‘species’ are defined at the operational taxonomic unit (OTU) level. After the pre-processing steps, the number of samples *m* varies between 145 and 195, and the number of species *N* varies between 387 and 1061 across the different body sites (see Methods). After the pre-processing stage, we apply the kNN algorithm described above. The prediction error calculation is performed in the following way. For each dataset, we divide the data into two parts. The first, containing 99% of the data is used as the training set, and the other 1% (typically 2 samples) serves as the test samples. For each test sample, we apply the kNN algorithm using different *k* values and calculate the prediction error between the predicted and the real abundance profiles. We repeat this process 1000 times, each time randomly dividing the training set into 99%/1% training/test samples, and calculate the errors. The average error versus *k* is presented as the blue line in Fig. 3**a-m**. The prediction error associated with the null model, a constant value corresponding to *k* = *m* is marked by the horizontal orange line.

**Figure 3.**
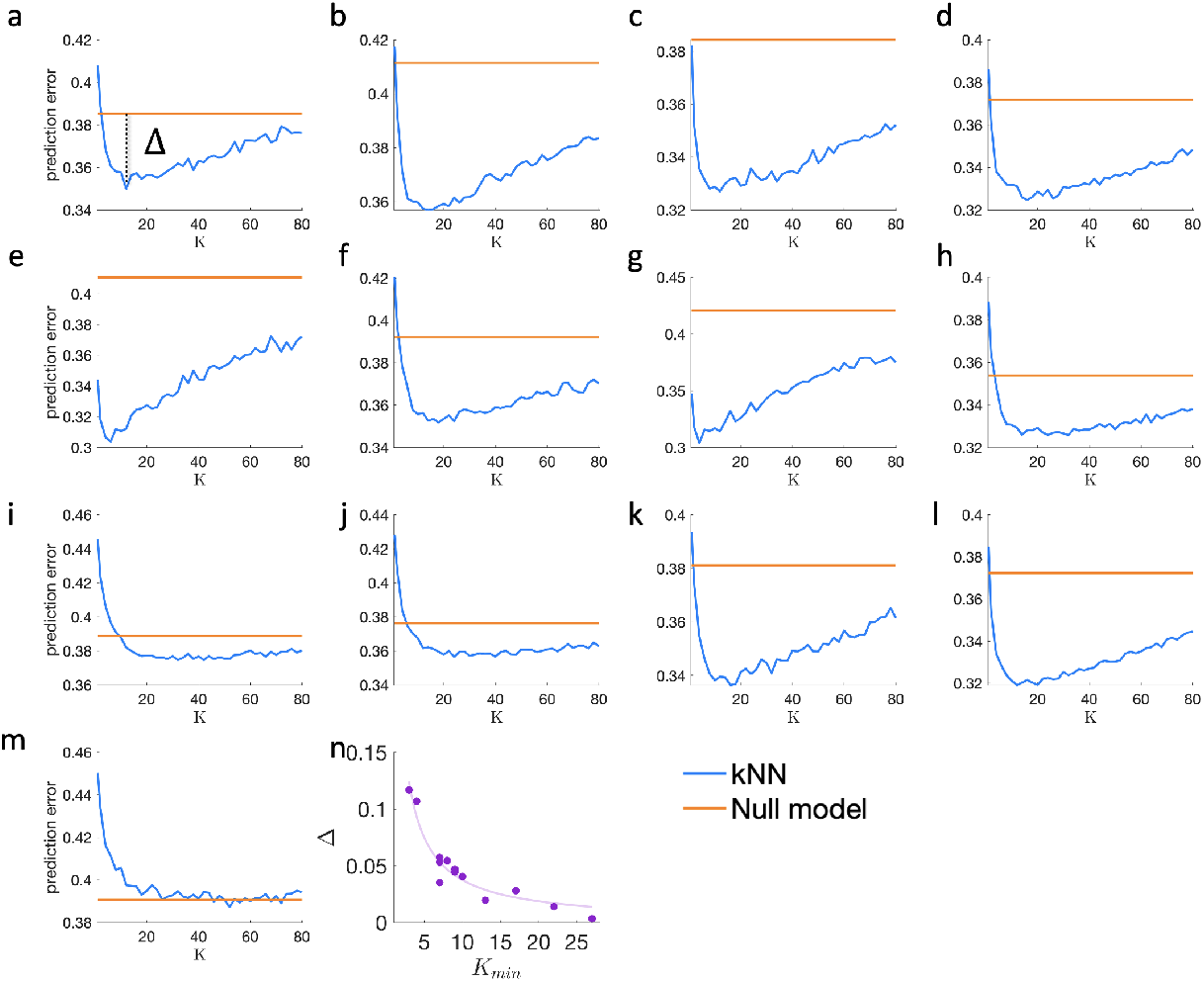
Predicting microbial composition in real metagenomic data. The prediction error of the kNN versus *k* (blue lines), compared with the prediction error of the null model(orange lines). Each point represents the average prediction error over 100 ‘leave-one-out’ realizations, where a single microbial sample is randomly selected as a test sample. Different panels show the results of 13 microbial communities from different body sites, obtained from the HMP dataset: anterior nares **(a)**, attached keratinized nares **(b)**, buccal mucosa **(c)**, hard palate **(d)**, left retroauricular crease **(e)**, right retroauricular crease **(f)**, palatine tonsils **(g)**, saliva **(h)**, subgingival plaque **(i)**, supragingival plaque **(j)**, throat **(k)**, tongue dorsal **(l)**, and stool **(m)**. In each case, we define Δ as the difference between the prediction errors of the kNN for *k* = *k*_*min*_ and of the null model, as demonstrated in **a. n**, Purple dots represent the relation between Δ and *k*_*min*_ calculated for the different body sites. The solid curve represents a polynomial fit.

We found that for small values of *k*, the prediction error decreases, reaching a minimum point *k*_*min*_, and then increases for larger values of *k*. We define the kNN gain, Δ, which quantifies the difference between the null model’s prediction error and the kNN’s prediction error for *k* = *k*_*min*_, as illustrated in Fig. 3**a**. The different body sites differ both in the values of *k*_*min*_ and of Δ, where the value of Δ is typically large when *k*_*min*_ is small and vice versa, as shown in Fig. 3**n**. As mentioned above, the value of *k*_*min*_ reconciles between specificity, i.e., averaging over the closest neighbors, and improving the statistics by averaging over a large number of samples. Accordingly, when *k*_*min*_ is relatively small, the closest vicinity of the test sample provides enough information for obtaining significantly better predictions than the null model, and, thus, a larger value of Δ. In contrast, a high value of *k*_*min*_ means that more samples are required and, thus, the kNN predictions are getting closer to the null model, i.e., a smaller value of Δ.

The key assumption in applying the kNN method for predicting microbial communities is that samples with similar species assemblages are expected to have similar abundance profiles. To investigate how the validity of this assumption across datasets is related to the performance of the kNN, we utilize the dissimilarity-overlap curve (DOC) analysis, a recently developed approach for the analysis of the relationships between species assemblages (measured as overlap) and abundance profiles (measured as dissimilarity) [16] (see Methods). The DOC represents the dissimilarity and overlap values calculated between a pair of microbial samples as a point in the dissimilarity-overlap plane. Typical examples of DOC clouds representing all sample pairs in a given dataset are shown in Fig. 4**a-c** for simulated data. If samples with similar species assemblages (high overlap) tend to have similar abundance profiles (low dissimilarity), the DOC cloud will have a negative slope at the range of high overlap (Fig. 4**a**). Thus, we can relate to the DOC slope at the range of high overlap as a proxy for the validity of the kNN assumption mentioned above.

**Figure 4.**
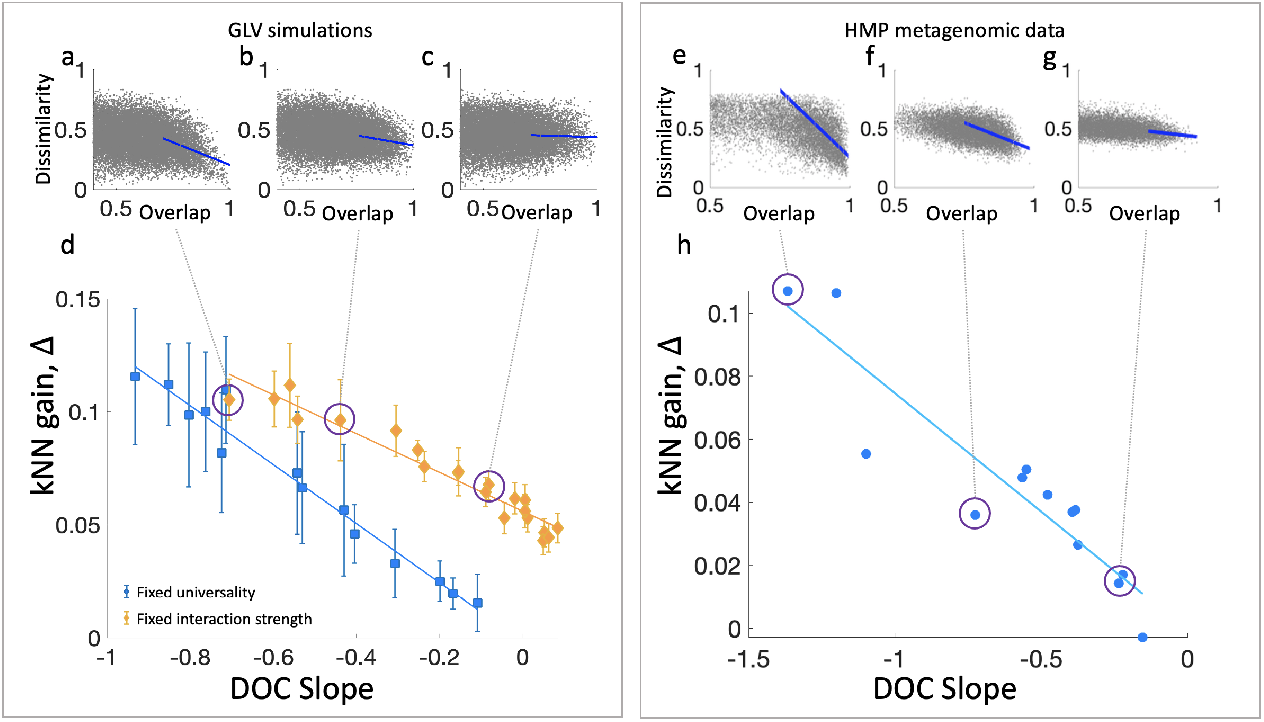
Dissimilarity-overlap analysis of the training data as a proxy for the kNN performance. **a-c**, Examples of dissimilarity-overlap curves (DOCs) for three cohorts of *m* = 1000 simulated samples, with *N* = 40 species, interaction strength *σ* = 3.4, and different universality values (λ = 0, 0.25, 0.5). The slope of the DOC is calculated using linear fit over the highest 20% overlapping points (blue lines). **d**, The kNN gain, Δ, versus the DOC slope for different cohorts of simulated samples. For each cohort, we calculate its DOC slope as well as its Δ value, calculated based on 50 test samples. Different cohorts were generated by choosing the ‘interaction strength’ and the ‘universality’ features. Blue symbols represent the mean Δ over 10 cohorts with a fixed universality value of λ = 0 and interaction strength #x03C3; ranging between 0.8 and 3.4. Yellow diamonds represent the mean Δ over 10 cohorts with a fixed interaction strength #x03C3; = 3.4 and λ between 0 and 1. The error bars represent the standard error (SE). The straight lines are linear fits (with goodness of fit *R*^2^ = 0.99 for both lines). **e-g**, Examples of DOCs of three cohorts of real microbial samples from different body sites (‘left retroauricular crease’, ‘hard palate’, and ‘subgingival plaque’). **h**, Same as **d**, for 13 body sites of the HMP dataset. The straight line is a linear fit (goodness of fit *R*^2^ = 0.96). In both simulated and real microbial data, the kNN gain is larger for cohorts with steeper DOC slopes (larger absolute values).

Here, we compare different datasets of both simulated and real microbial samples, and investigate the effect of different DOC slopes on the kNN predictions. We first generate cohorts of simulated samples with different DOC slopes. As demonstrated in Ref. [16], the slope of the DOC in GLV simulations can be controlled by tuning the ‘interaction strength’ or the ‘universality’ features of the underlying GLV models (see Methods). For each cohort, we calculate two quantities: i) The DOC slope, defined as the incline of the linear fit of 20% of the points with the highest overlap; ii) The kNN gain, Δ. Figure 4 **d** shows that Δ is correlated with the slope of the DOC such that better kNN predictions (higher Δ) are related to steeper DOC slopes (higher absolute values of the slopes). This pattern repeats for both tuning methods, i.e., changing the universality with a fixed interaction strength or vice versa. We then repeat the same analysis for the real metagenomic data analyzed in Fig. 3, measuring the DOC slope and the Δ between the kNN and the null model for each cohort. As shown in Fig. 4**h**, the kNN predictions are more accurate (larger Δ) for cohorts with steeper DOC slope. These results display a strong dependence between the DOC slope and the kNN predictability, supporting the relationship between the kNN assumption and its performance. Consequently, the DOC analysis can be used as a pre-analysis indicator of the suitability of the kNN method for a particular system.

Up to this point, we’ve assessed the mean kNN gain across a set of samples originating from a shared ecological environment. While utilizing the mean is logical when dealing with uniform groups, certain microbial communities might exhibit greater heterogeneity in the density of microbial samples. Since the kNN focuses on the neighboring samples, its performance may vary notably across different individuals in the same cohort.

In particular, the gut microbiome is a complex ecosystem with a considerably large number of detected species, and in the HMP dataset, its cohort showcases the largest sample-to-sample variability (the average Jaccard distance is 0.86 compared with between 0.78 and 0.84 in the other body sites). Figure 5**a** visualizes the ‘assemblage space’ of the gut microbiome (Principal coordination analysis using the Jaccard distance), where some areas of the assemblage space are denser than others. For example, the sample-to-sample distances between the subset of 15 samples marked by the green rectangle are significantly smaller compared with those of the 15 samples marked by the red rectangle (see inset).

**Figure 5.**
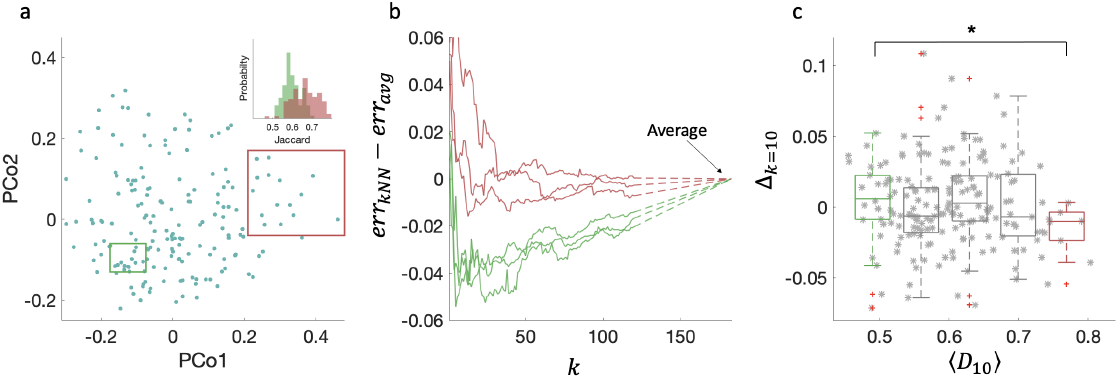
Individualized kNN analysis for the gut microbiome. **a**, The distribution of 184 microbial stool samples over the ‘assemblage space’, which represents the Jaccard distance between the species assemblages as depicted through principal coordinates analysis, is non-homogeneous. For example, the Jaccard distances between the 15 points marked by the red rectangle are significantly larger compared with the distances between the 15 points marked by the green rectangle (inset). **b**, Three examples of kNN profiles of samples with small values of ⟨*D*_(10)_⟩, i.e., the average Jaccard between the sample and its 10 nearest neighbors (0.45, 0.45 and 0.48, green curves) and three with high values of ⟨*D*_(10)_⟩ (0.74, 0.74 and 0.8, red curves). The kNN profiles of the samples from dense neighborhoods typically reach a minimum (best prediction) at small values of *k*. In contrast, the kNN predictions of samples from sparse neighborhoods typically require including many or even all the cohort samples. **c**, The kNN gain Δ of a predetermined value *k* = 10 is significantly higher for the samples with small ⟨*D*_(10)_⟩ values compared with those with large ⟨*D*_(10)_ ⟩ values. Box plots represent the first and third quartiles; middle line, median; red daggers, outliers. Boxes represent equal ranges of ⟨*D*_10_⟩ values. Asterisk, p value= 0.02, calculated using the Mann-Whitney U-test between the samples associated with the left and the right boxes.

We evaluate the kNN method for each individual and test whether it relates to the density of available samples in its local surroundings in the assemblage space. Specifically, we measure for each sample the average Jaccard distance between its species assemblage and its 10 nearest neighbors, ⟨*D*_10_⟩. Figure 5**b** demonstrates that the kNN profile, i.e., the kNN gain versus *k*, depends on the value of ⟨*D*⟩ for the analyzed individual. The kNN profiles of samples from dense areas in the assemblage space (small values of ⟨*D*⟩) have a typical minimum point at *k ≈*10 −20. This means that for these samples, the kNN prediction when using a small number of neighbor samples is better than the prediction that incorporates all the cohort’s samples (*err*_*kNN*_ < *err*_*avg*_). In contrast, the kNN profiles of samples from sparse areas in the assemblage space have shallower minimum points at large values of *k*, i.e., the prediction using the average is typically better than the kNN predictions using their closest neighbors.

In order to elucidate the efficacy of the kNN algorithm within the context of perturbation experiments, we conducted a comprehensive analysis on the microbiome datasets originally presented in Ref. [25]. This particular dataset encapsulates microbiome profiles for a cohort of 21 human subjects, both prior to and subsequent to the administration of antibiotics. Specifically, for each subject, two distinct samples were assessed: an initial sample, denoted as ‘baseline,’ collected prior to antibiotic treatment, and a subsequent sample, labeled ‘post-ABX,’ obtained at the 56-day mark following antibiotic administration (refer to the Methods section for details).

We utilize the kNN method to estimate the ‘post-ABX’ abundance profiles for each participating subject. Importantly, we aimed to scrutinize the impact of sample density within the assemblage space on the performance of the kNN algorithm. Given the heterogeneous distribution of post-ABX assemblages in this space — that is, some samples reside in regions characterized by higher sample density compared to others (Fig. 6a) — we ask whether the efficacy of the kNN predictions is associated with the local density of samples within their proximate neighborhoods. For each subject, we used a training set that comprised samples from all other subjects, while intentionally excluding the ‘baseline’ sample from the individual under prediction. This exclusion was executed to mitigate the risk of ‘data leakage,’ which could otherwise introduce bias into the model by capturing the ‘recovery effect’—a phenomenon wherein the post-antibiotic microbiome tends to revert towards its original state. In addition, while some of the subjects underwent spontaneous recovery, others were treated with probiotics or auto fecal microbiota transplant, which degraded or improved their recovery (see Ref. [26]). By excluding the ‘baseline’ sample from each subject, these differences are ignored and in our results, we didn’t see a significant difference between the groups.

**Figure 6.**
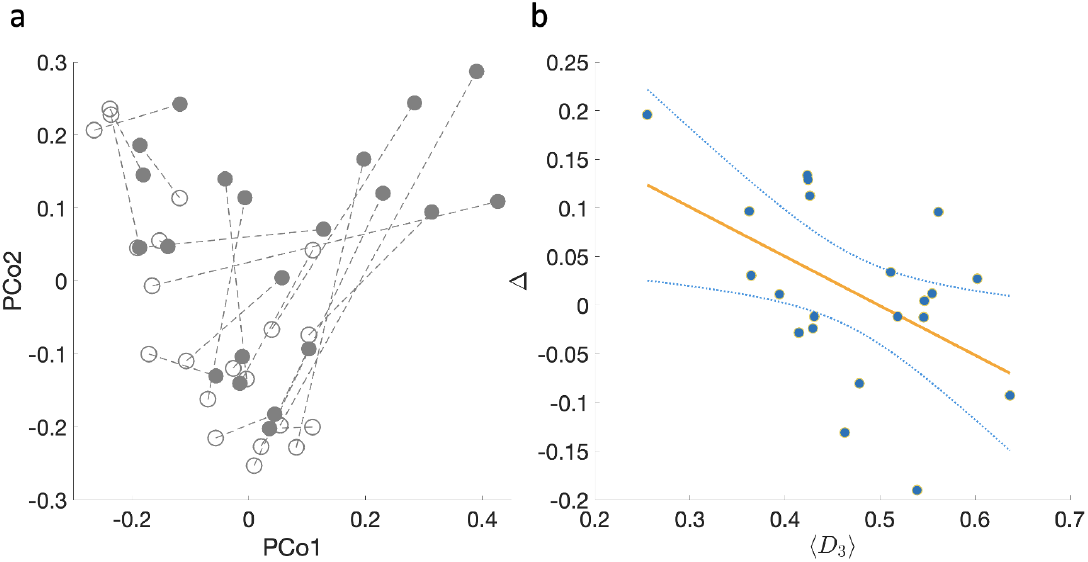
Association between kNN efficacy and local samples’ density in post-perturbation experiments. **a**, Principal coordination analysis of the Jaccard distances between 42 samples from 21 subjects, before (hollow circles) and after antibiotic administration (filled circles). Dashed lines are drawn between samples from the same subject. **b**, The kNN gain Δ versus the mean distance of the closest three neighbor samples ⟨*D*_3_⟩ calculated for 21 post-antibiotic microbial samples. Samples in denser regions in the assemblage space tend to have better predictions of the kNN method compared with samples in sparse regions. The orange straight line represents the linear regression model, represented by the equation *Y* = −0.5*X* + 0.25, with p-value=0.02. The light-blue curves indicate the 95% confidence envelop around the regression line.

Figure (Fig. 6b) shows that the kNN success (high value of Δ) is associated with narrower local neighborhoods of the predicted samples (lower value of ⟨*D*_3_⟩) (Pearson correlation −0.5, p-value 0.02). This outcome illustrates the implementation of the kNN method for forecasting the abundance profile of microbial communities following perturbations. As elaborated in the subsequent Discussion section, kNN can be effectively employed in conjunction with a predictive model for species assemblage or to facilitate the design of specific perturbations needed to guide the microbiome toward a desired stable state.

## Discussion

In this study, we predict species’ abundance profiles based on their presence/absence configuration using the kNN regression algorithm. The uniqueness of this approach lies in the fact that the kNN algorithm does not train a predictive model (such as a neural network model), but relies instead on comparing each test sample to the training data directly. Generally, avoiding the daunting task of accurately training a model has the advantages of saving precious time and computational resources, as well as reducing the risk of overfitting the model to a particular ecological environment. On the other hand, using the kNN method requires storing the whole data at the prediction step, unlike a trained model, which is much more compact. The priority that is given to one consideration over another depends on the ratio between the amount of data available and the complexity of the system. In cases where the system is relatively simple, e.g., an ecosystem with a small number of interacting species, and a large number of training samples, an effective predictive model can be accurately trained. However, natural microbial ecosystems typically consist of hundreds of species, while only a few dozen samples are available. In this case, the kNN approach might be an effective tool for the task of predicting the microbiome composition.

The essence of the approach we take in this research is to divide the complicated problem of predicting perturbation outcomes into a two-step prediction scheme: predicting the species assemblage, and - based on the assemblage- predicting the associated abundance profile. In practice, the two-step approach can be used in two main avenues: i) Predicting the response of a given perturbation using the kNN depends on the prior prediction of the assemblage. In this sense, while the kNN is “model-free”, the entire process does depend on a predictive model for the species assemblage. ii) Conversely, our strategy can be adapted to create a custom perturbation, with the goal of guiding a given subject’s microbiome towards a specified state. Here, the kNN is pivotal in selecting the target species assemblage that aligns with the intended abundance profile, providing direction for the intervention.

We have established that the kNN algorithm, when applied to cross-sectional data, yields the best results when using a relatively small number of neighboring samples. Typically, the value of *k*_*min*_ is about 10, which is only around 5 percent of the total available data. The fact that the algorithm uses only the nearest neighbors to predict the test sample most effectively is consistent with the hypothesis that microbial samples represent different alternative steady states of the ecosystem. We consider each microbial sample as a representation of a possible steady state of the ecosystem, whereas temporal changes are considered as minor fluctuations around a steady state, or transitions between alternative steady states, as also suggested by previous longitudinal studies [27, 28] and a macroecological description of the microbiome [29].

Accordingly, in order to make an accurate assessment of the test sample, the most relevant samples are those that are near its steady state, rather than the entire spectrum of alternative states. Lastly, we show that the kNN method’s main prerequisite is a relationship between the species assemblages of the samples and their abundance profiles, which can be characterized by the DOC analysis as the slope at a high overlap region. For this sake, we do not have to assume any interpretation regarding what factors lead to the variance of DOC slopes across different environments. Nevertheless, the feature that we used in our GLV simulations may suggest two qualitative ecological features of the data that affect the kNN’s performance. A relationship between species assemblages of different samples and their abundance profiles may be a result of host-independent underlying mechanisms that control the species dynamics, i.e., consistent ecological environment-species and species-species interactions across different hosts in the same ecological environment. Accordingly, an ecological environment for which the kNN can be successfully applied may (but not necessarily) be characterized by strong species-species interactions and ‘universal’ underlying dynamics.

## Methods

### Population dynamics model

The GLV model represents the dynamics of *N* interacting species as a set of ordinary differential equations: 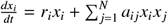. Here, *r*_*i*_ is the intrinsic growth rate of species *i, a*_*i j*_ is the interaction strength between species *j* and *i*, and 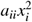 (with *a*_*ii*_ < 0) represents the logistic term. We consider a microbial ‘sample’ as a steady state of a GLV model parameterized by the growth rate vector **r** = {*r*_*i*_} ∈ ℝ^*N*^ and the interaction matrix *A* = (*a*_*i j*_) ∈ ℝ^*N*×*N*^. In our simulations, *r*_*i*_ is randomly chosen from the uniform distribution 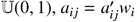, where 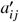 is randomly chosen from the normal distribution ℕ(0, #x03C3;) with interaction strength #x03C3; = 0.6 for *N* = 10, 20, 40 and #x03C3; = 0.4 for *N* = 100 (to ensure system’s stability) and *w*_*i*_ weighs the influence of species *i* (*w*_*i*_ is chosen from a power-law distribution such that 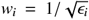 where *ϵ*_*i*_ is randomly chosen from the uniform distribution 𝕌(0, 1) and then *w*_*i*_ is normalized such that 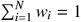, and we set *a*_*ii*_ = −1. We generated different ‘cohorts’, each consisting of *m* ‘samples’ (unique steady states), generated by integrating the GLV differential equations with random initial conditions (both initial assemblage and abundance profile were randomly chosen). When tuning the ‘universality’ feature, we use for each cohort GLV models that differ from each other in their specific parameters. For each cohort, we first construct a ‘base’ GLV model (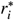 and 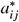) as before, and for each realization, we create 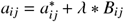, where *B*_*i j*_ is randomly chosen from a normal distribution ℕ(0, 1).

### Overlap between species assemblages

Given two microbial samples, represented by two abundance vectors *x* = (*x*_1_, *x*_2_, …, *x*_*n*_) ∈ ℝ^*N*^ and *y* = (*y*_1_, *y*_2_, …, *y*_*n*_) ∈ ℝ^*N*^, their species assemblages are denoted as *X* = {*i*| *x*_*i*_ > 0} and *Y* = {*i*| *y*_*i*_ > 0}.

To quantify the similarity of the species assemblages of the two samples, we use the overlap measure 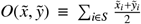 where 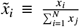 and 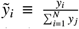 are the relative abundances, and *S* ≡ *X* ⋂ *Y* is the set of the shared species present in both samples. If *S* is empty, 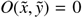. In the case of {*S* = 1, 2, …, *N*}, that is, all the species in *X* and *Y* are shared, 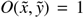, but the abundances can be different.

### Dissimilarity between abundance profiles

To compare the abundance profiles of two samples, we first renormalize the relative abundances of only the shared species (in set S), yielding 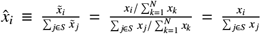 and *ŷ*_*i*_ is defined similarly. By using this definition, we remove the spurious dependence between the relative abundances of the shared and the non-shared species. More importantly, this renormalization assures that the calculated dissimilarity measure is mathematically independent of the overlap measure. The dissimilarity is then calculated via the root Jensen-Shannon divergence (rJSD) which is defined by 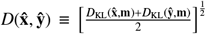 where 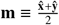 and 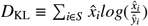 is the Kullback-Leibler divergence between 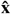 and **ŷ**.

### DOC

To systematically compare sample pairs with a wide range of overlap values and analyse their dissimilarity–overlap relations, we calculate the overlap and dissimilarity of all the sample pairs from a given set of microbiome samples, and represent each sample pair as a point in the dissimilarity–overlap plane. The DOC slope is calculated using linear fit over the points with the top 20% overlap values using MATLAB’s `polyfit` function.

### Jaccard similarity

The Jaccard coefficient of two sets, *A* and *B*, is defined as the size of the intersection divided by the size of the union of the sample sets 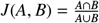. If the two sets *A* and *B* are equal, *J*(*A, B*) = 1, and if *A* and *B* are disjoint sets, *J*(*A, B*) = 0.

### Null model

For a training set Θ of *m* samples representing the abundance profiles of *N* species, the abundance profile of the test sample Φ is predicted to be 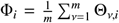 where ϕ_*i*_ = 1, and Φ_*i*_ = 0 where ϕ_*i*_ = 0 (The null model is similar to the kNN where *k* = *m*).

### Machine Learning

We train fully connected neural networks with two hidden layers, where each layer consists of 50 neurons, and apply the Levenberg–Marquardt backpropagation algorithm for optimization. The Number of neurons in the first and last layers is equal to the number of species *N*. Each cohort of samples is first divided into training, validation and test sets, consisting of 70%, 20%, 10% of the samples, respectively. We use the species assemblage of each sample as the input, and the relative abundance as the desired output when training the network. The loss is calculated using the mean square error (MSE) between the real and predicted abundance profiles.

### Data and data pre-processing

We use cross-sectional microbiome data from the Human Microbiome Project (HMP)[1], a 16S rRNA gene-based dataset of the human microbiomes from 239 healthy subjects. We performed the analysis at the OTU level. The cohorts of samples are pre-processed as follows: We remove from each cohort low-abundance OTUs with less than one read per sample on average, across all the samples in the cohort. Additionally, we remove OTUs with a prevalence lower than 10% in each cohort. We used a single sample from each subject. In cases where more than one sample was available, we used the first collection. After the pre-processing stage, the cohorts contain the following numbers of samples and species (*m* is the number of samples and *N* is the maximal number of species, respectively). Anterior nares (*m* = 145, *N* = 497), attached keratinized nares (*m* = 195, *N* = 585), buccal mucosa (*m* = 179, *N* = 570), hard palate (*m* = 172, *N* = 672), left retroauricular crease (*m* = 159, *N* = 387), palatine tonsils (*m* = 181, *N* = 850), right retroauricular crease (*m* = 163, *N* = 403), saliva (*m* = 163, *N* = 804), subgingival plaque (*m* = 179, *N* = 942), supragingival plaque (*m* = 183, *N* = 897), throat (*m* = 172, *N* = 759), tongue dorsum (*m* = 184, *N* = 803), and stool (*m* = 184, *N* = 1061). Full protocol details are available at the HMP DACC website (http://hmpdacc.org/HMMCP).

## Availability of data and materials

Data from the Human Microbiome Project can be downloaded at https://hmpdacc.org/. The code for main methods used in this project is available at https://github.com/eitanas/Model-free-Prediction-of-Microbiome-Compositions.

## Notes

### Competing Interest Statement

The authors have declared no competing interest.

### Summary of Updates

Personalized assessment of the kNN: We examined the performance of the kNN method for predicting microbial abundances not only for a group of samples but also for an individual sample, finding that this is affected by the density of samples in the local environment. Demonstration of the kNN in perturbation experiments: We analyzed microbial samples from experiments of human subjects before and after antibiotic administration, showing that the kNN algorithm is effective in predicting the abundance profiles of the perturbed microbiomes. We have added Fig.5 and Fig.6 which describe the results on the new data.

